# Characterization of imprinted genes in rice reveals post-fertilization regulation and conservation at some loci of imprinting in plant species

**DOI:** 10.1101/143214

**Authors:** Chen Chen, Tingting Li, Shan Zhu, Zehou Liu, Zhenyuan Shi, Xiaoming Zheng, Rui Chen, Jianfeng Huang, Yi Shen, Shiyou Luo, Lei Wang, Qiao-Quan Liu, E Zhiguo

## Abstract

Genomic imprinting is an epigenetic phenomenon by which certain genes display monoallelic expression in a parent-of-origin-dependent manner. Hundreds of imprinted genes have been identified from several plant species. Here we identified, with a high level of confidence, 208 imprinted candidates from rice. Imprinted genes of rice showed limited association to the transposable elements, which is contrast to the findings in *Arabidopsis*. Generally, imprinting of rice is conserved within species, but intraspecific variations were confirmed here. Imprinting between cultivated rice and wild rice are likely similar. The imprinted genes of rice do not show significant selective signatures overall, which suggests that domestication imposes limited evolutionary effects on genomic imprinting of rice. Though the conservation of imprinting in plants is limited, here we prove that some loci tend to be imprinted in different species. In addition, our results suggest that differential epigenetic regulation between parental alleles can be established either prior to or post-fertilization. The imprinted 24-nt small RNAs, but not the 21-nt ones, likely involve the regulation of imprinting in an opposite parental-allele targeting manner. Together, our findings suggest that regulation of imprinting can be very diverse, and genomic imprinting as well as imprinted genes have essential evolutionary and biological significance.

## Introduction

Genomic imprinting has been observed in many species, from mammals to flowering plants (Pires and Grossniklaus, 2014). The imprinted genes only express one of the parental alleles in a parent-of-origin-dependent manner (Köhler et al., 2012; Gehring, 2013). Genes that exclusively or preferentially express the maternal or paternal alleles are termed maternally expressed genes (MEGs) or paternally expressed genes (PEGs), respectively. Hundreds of putative imprinted loci, including protein-coding genes and non-coding RNAs, have been discovered in plant (Gehring et al., 2011; Hsieh et al., 2011; Luo et al., 2011; Waters et al., 2011; Wolff et al., 2011; Zhang et al., 2011; Waters et al., 2013; Xin et al., 2013; Pignatta et al., 2014; Xu et al., 2014; Florez-Rueda et al., 2016; Hatorangan et al., 2016; Klosinska et al., 2016; Zhang et al., 2016). The conservation of imprinting between plant species is very limited (Waters et al., 2013; Hatorangan et al., 2016), which indicates that the evolution of genomic imprinting in plants is very rapid. Genomic imprinting is epigenetically regulated by either DNA methylation or histone modification, and in some circumstances, both mechanisms are involved (Huh et al., 2008; Köhler and Weinhofer-Molisch, 2010). In plants, imprinting is predominately expressed in endosperm (Luo et al., 2011; Raissig et al., 2013; Klosinska et al., 2016). Overall, the endosperm is hypomethylated in comparison with the embryo and vegetative tissues (Gehring et al., 2009; Zemach et al., 2010; Rodrigues et al., 2013; Xing et al., 2015; Klosinska et al., 2016). DNA glycosylase *DEMETER* (*DME*), which is responsible for DNA demethylation, is predominantly expressed in central cell before fertilization (Choi et al., 2004). Imprinting disorders can be found in *dme* mutants (Hsieh et al., 2011; Wolff et al., 2011; Vu et al., 2013), indicating that proper DNA methylation status is required for the establishment of genetic imprinting in the endosperm. Several studies revealed that imprinted genes usually neighbor transposable elements (TEs) (Wolff et al., 2011; Pignatta et al., 2014; Hatorangan et al., 2016). High expression of *DME* in the central cell promotes demethylation of TEs and their adjacent imprinted genes (Gehring et al., 2009). Therefore, the maternal alleles of imprinted genes are activated in the central cell, but the paternal alleles keep silenced in sperm cell due to hypermethylation, which eventually leads to maternal monoallelic expression after fertilization. Polycomb Repressive Complex 2 (PRC2)-mediated Histone H3 Lysine 27 Trimethylation (H3K27me3) is required for proper genomic imprinting (Huh et al., 2008; Köhler and Weinhofer-Molisch, 2010). Mutation of *fertilization-independent endosperm (FIE)*, a member of FIS-PRC2, may result in disruption of imprinting at some loci (Hsieh et al., 2011; Wolff et al., 2011).

Genetic evidence showed that several MEGs are likely very important for seed and/or endosperm development in *Arabidopsis* (Chaudhury et al., 1997; Luo et al., 2000; Tiwari et al., 2008; Gerald et al., 2009; Costa et al., 2012; Liu et al., 2014). However, many mutants of confirmed imprinted genes, usually PEGs, show no phenotypes (Köhler et al., 2005; Shirzadi et al., 2011; Bratzel et al., 2012; Vu et al., 2013; Wolff et al., 2015). In addition, imprinting conservation between phylogenetically close species is limited (Waters et al., 2013; Hatorangan et al., 2016). These observations have cast doubt on the importance of genomic imprinting in plants. The kinship theory (Haig and Westoby, 1991) hypothesizes that imprinting arose as a consequence of a conflict of interest between the male and female gametes, according to which the male will gain the benefit of maximal transmission of resources from the mother to the offspring, whereas the female benefits by suppressing the offspring’s growth-related demands, which are driven by paternally active genes, to protect the mother. Many observations support the kinship hypothesis (Köhler et al., 2012; Pires and Grossniklaus, 2014; Rodrigues and Zilberman, 2015). However, in plants, this theory has been challenged by several controversies (Rodrigues and Zilberman, 2015). Recent findings in *Arabidopsis* revealed that some PEGs may be involved in hybrid incompatibility (Wolff et al., 2015), implying that genomic imprinting possibly plays a role in speciation. However, the evolutionary significance of genomic imprinting requires further investigation in different plant species.

In the present study, we extensively explored the genomic imprinting of rice. Our results showed that the regulatory mechanisms of genomic imprinting in rice might be very diverse. The findings will help us to better understand the regulation and evolution of imprinting in plants.

## Results

### Identification of imprinted genes in cultivated rice

Aiming to explore imprinted genes from rice, we used three-line hybrid rice strains to construct reciprocal crosses. A cytoplasmic male sterile (CMS) line carries a sterility gene encoded by the cytoplasmic genome, and the pollens it produces are completely sterile. A maintainer line shares the same nuclear genome as its corresponding CMS line but has distinct cytoplasmic genome. Due to the absence of a cytoplasmic sterility gene, a maintainer line is fertile. Using rice CMS and maintainer lines for crossing greatly increased the efficiency of hybridization and enabled us to rigorously control the timing window of fertilization, as well as to avoid false hybridization events. Here, we used the *japonica* CMS lines Liuqianxin-A and Yu6-A and their maintainer lines Liuqianxin-B and Yu6-B, the *indica* CMS lines Rongfeng-A and Wufeng-A and their corresponding maintainer lines Rongfeng-B and Wufeng-B to make two distinct reciprocal cross sets, namely Liuqianxin-A/Rongfeng-B (LR) and Rongfeng-A/Liuqianxin-B (RL), and Yu6-A/Wufeng-B (YW) and Wufeng-A/Yu6-B (WY). The LR and RL reciprocal crosses, and YW and WY reciprocal crosses are indicated as LR-RL and YW-WY hereafter for convenience. Meanwhile, we made crosses of Wufeng-A/Wufeng-B (WW), Rongfeng-A/Rongfeng-B (RR), Liuqianxin-A/Liuqianxin-B (LL), and Yu6-A/Yu6-B (YY), which resembled the selfing inbred lines, to be controls for the validation of imprinting. Morphological observations suggested that seeds produced by LR, RL, YW, and WY developed normally **(Figure 1A and B).** Endosperms at **5** days after fertilization (DAF) were collected from these reciprocal crosses for RNA-sequencing (RNA-seq). By genomic re-sequencing of Liuqianxin-A, Yu6-A, Rongfeng-B, and Wufeng-B, 62,102 and 59,107 SNPs, which corresponded to 12,299 and 11,044 endosperm expressed genes, were obtained to distinguish the parental alleles from the mRNA transcripts of LR-RL and YW-WY, respectively. In total, parental expression of 4,946 and 6,308 genes in LR-RL and YW-WY, respectively, showed deviation from the 2:1 maternal-to-paternal prediction by *χ*^2^ test (*p* < 0.05 and FDR < 0.05). Next, we surveyed parent-of-origin effects of these genes using the standard that the observed expression value should be 2-fold (for moderate imprinting) or 5-fold (for strong imprinting) higher than the expected value in both directions of the reciprocal crosses **(Figure 1C and D).** Therefore, for MEGs, the proportion of maternal alleles should be greater than 0.8 or 0.91, while for PEGs, it should be less than 0.5 or 0.29. According to this criterion, there were 177 and 118 MEGs identified above 0.8 and 0.91 levels from LR-RL, respectively. The corresponding numbers of MEGs were 347 and 148 in YW-WY hybrids **(Supplemental Table 1).** At the levels of 0.5 and 0.71, there were 504 and 237 PEGs identified from LR-LR and 362 and 129 PEGs identified from YW-WY **(Supplemental Table 1).**

**Figure 1.**
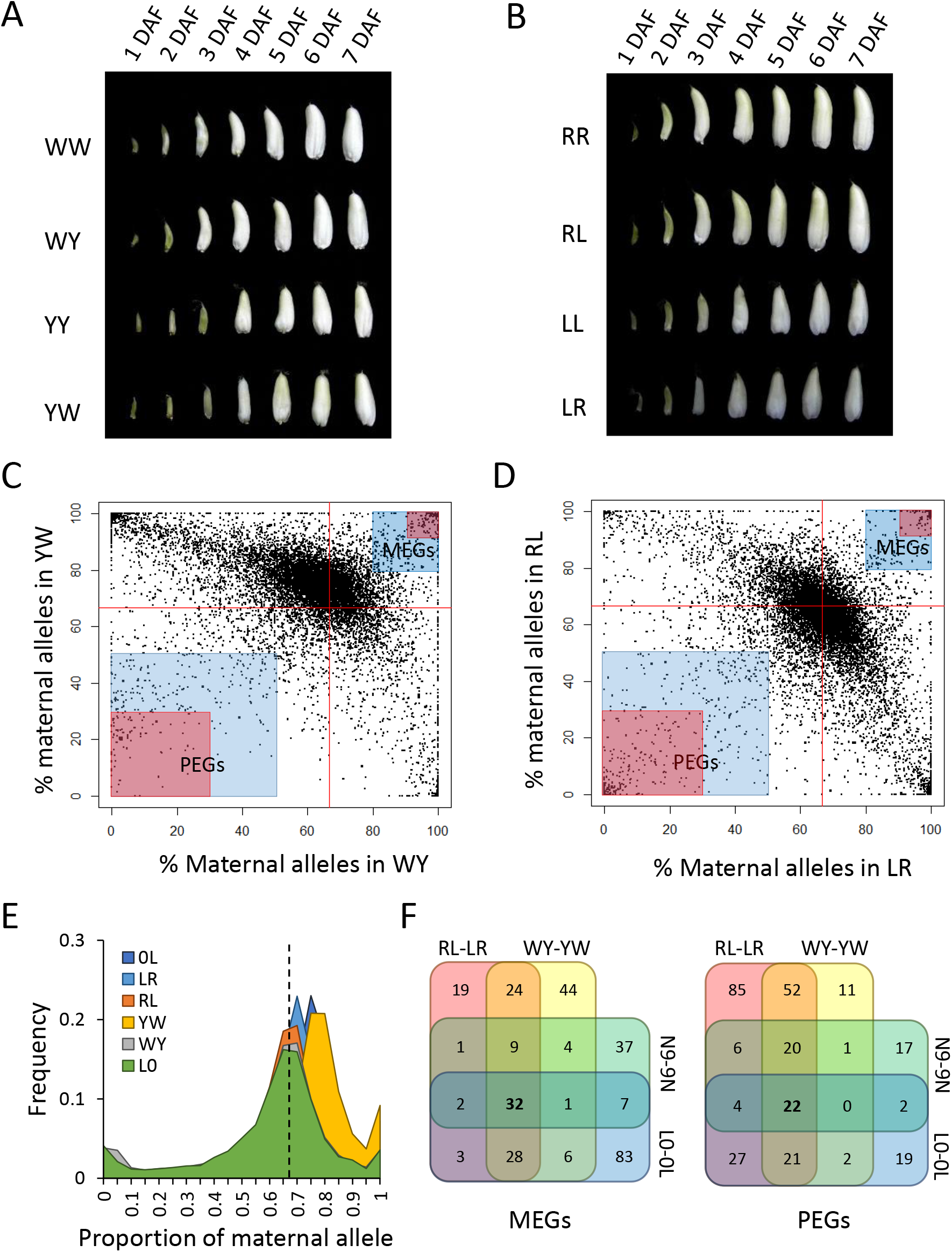
Identification of imprinted genes in rice. **(A)** Seed morphology of Wufeng-A x Wufeng-B (WW), Wufeng-A x Yu6-B (WY), Yu6-A x Yu6-B (YY), and Yu6-A x Wufeng-B (YW) from 1 day after fertilization (DAF) to 7 DAF. **(B)** Seed morphology of Rongfeng-A x Rongfeng-B (RR), Rongfeng-A x Liuqianxin-B (RL), Liuqianxin-A x Liuqianxin-B (LL), and Liuqianxin-A x Rongfeng-B (LR) from 1 DAF to 7 DAF. **(C, D)** Allele-specific expression analysis of WY-YW **(C)** and RL-LR **(D)**. The highlighted areas indicate moderate (2-fold higher the expected, blue) and strong (5-fold higher the expected, red) imprinted genes. Red lines indicate the 0.67 expect. The proportion of maternal alleles is calculated by FPKM_maternal_/(FPKM_maternal_+FPKM_paternal_). The dash line indicates the 0.67 expect. **(E)** Distribution of the proportion of maternal alleles at each loci with informative SNPs in WY, YW, RL, LR, Longtepu x 02428 (L0) and 0L. **(F)** Venn diagram of the imprinted genes identified from MY-YM, RL-LR, L0-0L and Nipponbare and 9311 reciprocal crosses (N9-9N).

In general, we identified more MEGs but less PEGs from YW-WY. By analyzing the ratio of (maternal transcripts): (maternal transcripts + paternal transcripts) at each locus, we found that the distribution peak of YW was shifted from the 0.67 to 0.75–0.8 **(Figure 1E),** which meant that globally, the mother Yu6-A contributed more mRNA transcripts than the expected 2:1 ratio. A similar trend was also observed in the cross of LR, albeit much less significant than that displayed in YW **(Figure 1E).** In contrast, most genes did not deviate from the expected ratio of 2:1 in WY and RL **(Figure 1E).** We believe that the deviation could not be caused by false hybridization events, because the CMS line Yu6-A is completely sterile and cannot, therefore, self-fertilize. Rice endosperm is easy to be separated from the maternal tissues at 5 DAF, and the two independent RNA-seq replications of YW showed high correlation (*r*^2^=0.9873), which led us to think that it was not likely the contamination of maternal tissues caused the deviation. We used the genomic sequence of Nipponbare, a *japonica* rice variety, as the reference; therefore, mapping bias could cause this deviation. Taking account of the fact that Rongfeng-B and Wufeng-B have a close relationship (http://ricedata.cn), but LR did not show significant deviation as YW, we supposed that it was low chance that the mapping bias caused the deviation. In light of the observations that *indica/indica* crosses (WW and RR) exhibited faster early endosperm development than the *japonica/japonica* crosses (LL and YY), and the hybrid seeds were more like their maternal parent **(Figure 1A and B),** we assumed that the deviation of maternal contribution at 5 DAF could be caused by slower endosperm development when *japonica* rice was used as the mother, which led to a slower maternal-to-zygotic transition. To confirm this hypothesis, we reanalyzed the RNA-seq data of the 7 DAF endosperm from the Longtepu (*indica*) and 02428 (*japonica*) reciprocal crosses (L0-0L)(Yuan et al., 2017). Consistent to our hypothesis, when the *japonica* rice 02428 used as the mother, the distribution peak of (maternal transcripts): (maternal transcripts + paternal transcripts) shifted to 0.75 **(Figure 1E).** These findings suggested that when *japonica* used as the mother, it contributes more for the early endosperm development of the *japonica/indica* hybrid. To minimalize the influence of the bias, in the following studies, we mainly focused on the 208 strong imprinted genes (>5-fold bias) shared between LR-RL and YW-WY for analysis **(Figure 1F).**

We conducted RT-PCR sequencing to validate the imprinting of 14 candidates. All the ones we tested were confirmed as imprinted genes **(Supplemental Figure 1A),** which convinced us that most of the imprinted candidates we identified are reliable. We also performed single-colony sequencing and pyro-sequencing **(Supplemental Figure 1B and C)** to unambiguously confirmed the imprinting status of *GW2* and *OsPPKL2*, two quantitative trait loci for rice seed development (Song et al., 2007; Zhang et al., 2012b), which implied that genome imprinting may directly function on seed development of rice.

Gene ontology (GO) analysis showed that for MEGs, the term “DNA binding” (*p* = 0.00031, FDR= 0.017) was enriched, whereas for PEGs, the terms “transferase activity” (*p*=6.3e-05, FDR=0.0014), “kinase activity” (*p* = 6.3e-05, FDR = 0.0014) and “protein binding” (p = 0.001, FDR = 0.015) were overrepresented. The result suggested that PEGs and MEGs might have different functions. Interestingly, we noticed that several of the imprinted genes are involved in epigenetic regulation **(Supplemental Table 2).** A previous study reported that *OsFIE1* is the only imprinted PRC2 gene in rice (Luo et al., 2009). Here we found that *OsEMF2a*, a Su(z)12 homolog of rice, is imprinted as well. Notably, the Su(z)12 member *FIS2* is also imprinted in *Arabidopsis* (Jullien et al., 2006).

### Comparison to previously identified imprinted genes in rice

Luo et al. (2011) conducted a genome-wide survey of imprinted genes in rice using the *japonica* cultivar Nipponbare and the *indica* cultivar 9311 as the parents (N9-9N). Approximately 74% (53/72) PEGs and 53% (49/93) MEGs they identified were overlapped with the ones identified in the present study **(Figure 1F).** Similarly, about 80% (76/97) PEGs and 44% (72/162) MEGs discovered from L0-0L (Yuan et al., 2017) were included in our candidates list **(Figure 1F),** nevertheless, overlaps between N9-9N and L0-0L were much less (Yuan et al., 2017). The results indicated that we discovered more candidates than previous studies, and many of the imprinted genes identified before were included in our candidates list. In total, 300 MEGs and 289 PEGs have been identified from rice so far **(Figure 1F and Supplemental Table** 1). It seemed that PEGs are more conserved than the MEGs in rice. About 54.3% PEGs (157/289) could be found in at least two sets of the four reciprocals (RL-LR, WY-YW, N9-9N and L0-0L), while only 39% MEGs (117/300) were coincided in different combinations.

Most of the imprinted genes identified in one combination also tend to be imprinted in other reciprocal crosses **(Figure 2A-C and Supplemental Figure 2).** The ones that only found in one set of the reciprocals usually lack informative SNPs or sufficient reads for the determination of imprinting in other crosses **(Supplemental Figure 2).** Allele-specific imprinting has been found in maize and *Arabidopsis* (Waters et al., 2013; Pignatta et al., 2014). Here, if an imprinted gene identified from one reciprocal-cross set failed to reach the criterion of moderate imprinting (<2-fold deviation from the 2:1 expect) in other reciprocal crosses, we considered it as an allele-specific imprinted gene. Seventy allelic specific imprinted genes were found in rice **(Supplemental Table 3).** For example, LOC_Os01g38650 was identified as an MEG in RL-LR, N9-9N and L0-0L **(Supplemental Table** 1). It was maternally expressed in YW, but became biallelically expressed in WY **(Figure 2D).** The imprinting variations of LOC_Os01g38650 and LOC_Os06g42790 were confirmed by RT-PCR sequencing **(Figure 2D).**

**Figure 2.**
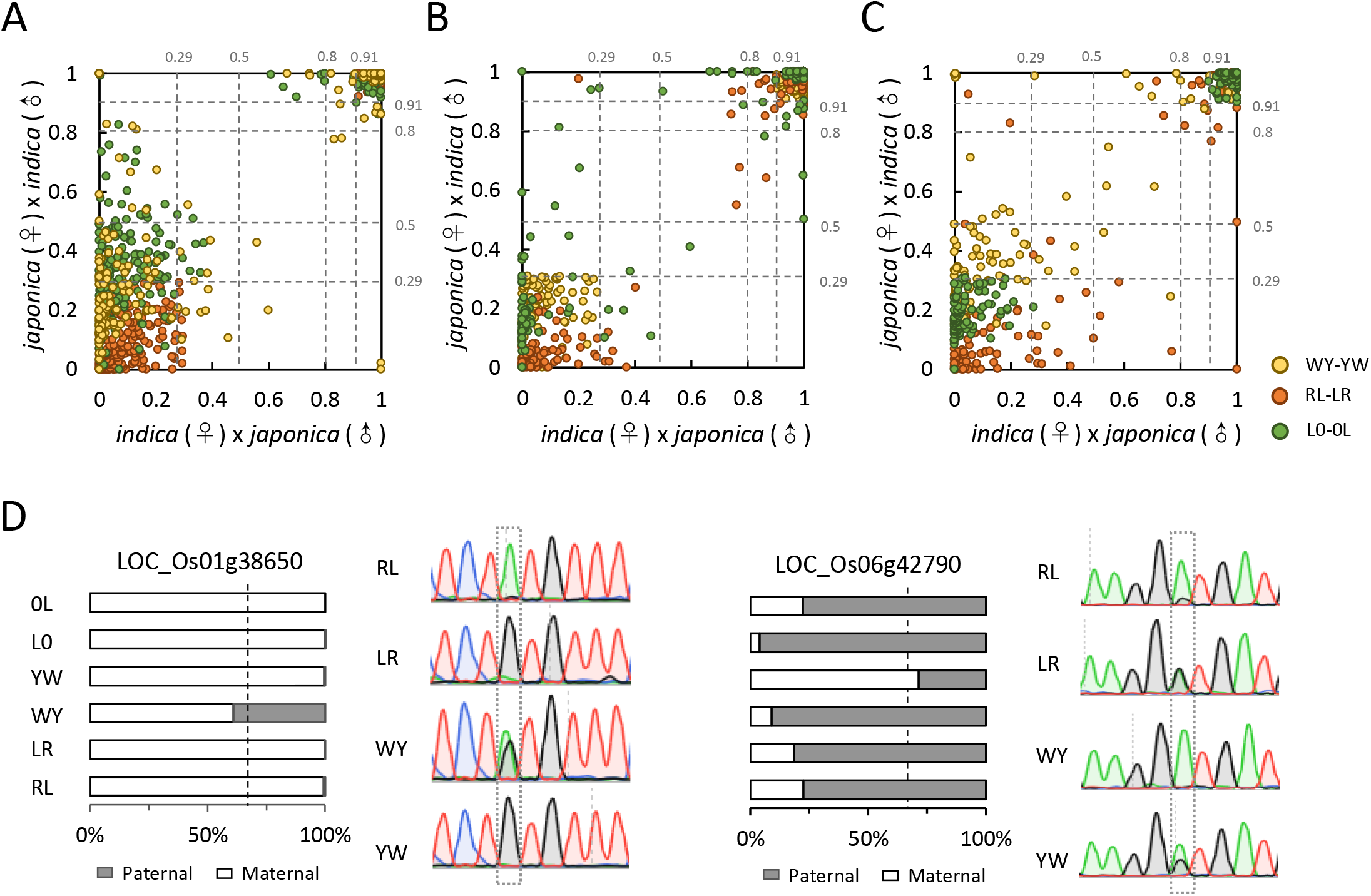
Intraspecific imprinting variation in rice. **(A-C)** Scatter diagrams of the proportion of maternal alleles (FPKM_matemal_/(FPKM_matemal_+FPKM_patemal_)) in *indica* (♀) × *japonica* (♂) and *japonica* (♀) × *indica* (♂) reciprocal crosses. **(A)** Proportion of maternal alleles of the imprinted genes identified from RL-LR in WY-YW and L0-0L; **(B)** Proportion of maternal alleles of the imprinted genes identified from WY-YW in RL-LR and L0-0L; **(C)** Proportion of maternal alleles of the imprinted genes identified from L0-0L in RL-LR and WY-YW. Most of the imprinted genes identified from one reciprocal-cross set are imprinted in other sets. **(D)** Examples of alleles-specific imprinting in rice. The bar charts show the proportion of parental alleles in different crosses; the sequencing diagrams show the validation of intraspecific imprinting variation by RT-PCR sequencing approach. Dash lines indicate the 0.67 expect, the informative SNPs are boxed by dash lines.

### PEGs of rice are less associated with TEs and tend to be clustered in genome

Transposable elements (TEs) are enriched in the proximity of imprinted genes in *Arabidopsis* and *C. rubella* (Wolff et al., 2011; Pignatta et al., 2014; Hatorangan et al., 2016). Yuan et al. (2017) found that miniature inverted-repeat transposable elements (MITEs) are enriched around the imprinted genes of rice. However, Luo et al. (2011) stated that the TE enrichment was not observed in rice. To investigate the controversy, we comprehensively searched the TEs in the rice genome. In total, 149,007 TEs were identified, which consist about 36% of the rice genome. When using the united imprinted genes of RL-LR, WY-YW, N9-9N and L0-0L for analysis, about 66% MEGs and 57% PEGs have a TE vicinity within the 2-kb flanking, while more than 70% genes encoded by the rice genome have a TE vicinity **(Figure 3A).** We did not find the enrichment of TEs within the 2-kb flanking regions of imprinted genes in rice. However, the chance of a PEG linking with a TE within the 2-kb flanking was lower than that of a MEG **(Figure 3A).** Consistent results were obtained when using different MEGs and PEGs that identified from distinct reciprocal crosses for analysis **(Figure 3A).** To minimize the influence of the TEs variation in different varieties, we used the 32 MEGs and 22 PEGs **(Supplemental Table 1)** that highly conserved among RL-LR, WY-YW, N9-9N and L0-0L for the analysis. The conserved PEGs still showed less association with TEs **(Figure 3A).** Statistical analysis showed that the distance of MEGs to their most close upstream and downstream TE vicinities was not different from the whole-genome control, but was smaller than that of the PEGs **(Figure 3B and C).** These data strongly suggested that the imprinted genes, especially the PEGs, showed limited association with TEs in rice.

**Figure 3.**
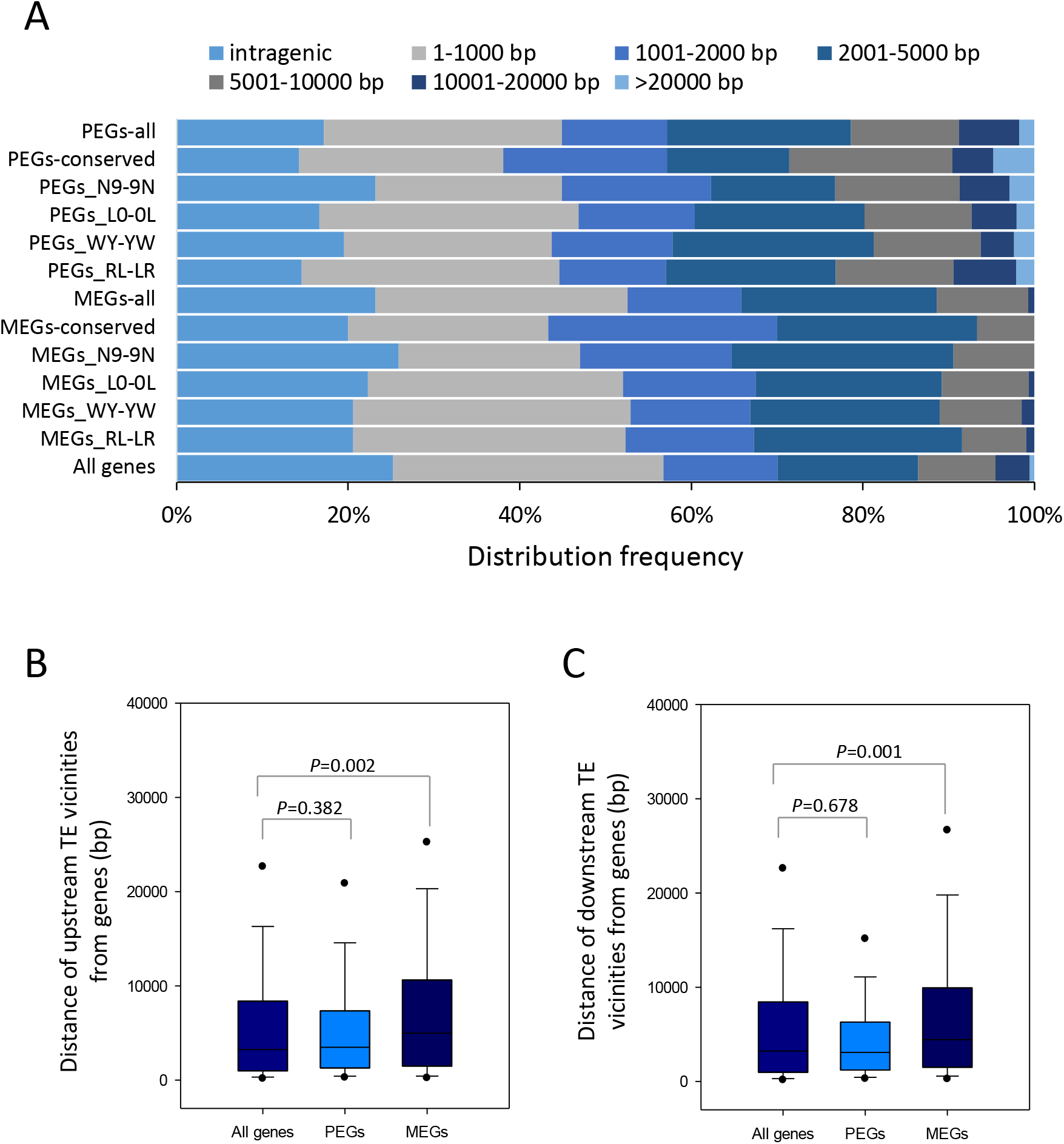
PEGs of rice are less associated with TEs. **(A)** Distribution frequency of the distance of rice imprinted genes to their TE vicinities. Neither MEGs or PEGs show TE enrichment within 2-kb flanking. PEGs-all and MEGs-all indicate the united PEGs and MEGs of RL-LR, WY-YW, N9-9N and L0-0L, respectively. PEGs-conserved and MEGs-conserved indicate the common PEGs and MEGs among RL-LR, WY-YW, N9-9N and L0-0L, respectively. **(B-C)** The united PEGs of rice has an overall larger distance to their upstream **(B)** and downstream **(C)** TE vicinities in comparison with the united MEGs; the overall distance of united MEGs to their upstream **(B)** and downstream **(C)** TE vicinities do not show significance to that of all the genes encoded by the rice genome. Kruskal-Wallis one way analysis is used for statistical analysis.

Mammalian imprinted genes are usually clustered in the genome (Edwards and Ferguson-Smith, 2007). However, this distribution pattern was not found in plants, though some micro-clusters were observed across the genomes of different plant species (Gehring et al., 2011; Luo et al., 2011; Wolff et al., 2011; Zhang et al., 2011). We used the united imprinted genes of MY-YM and RL-LR for clustering analysis. By a window sliding strategy, we found that 113 imprinted genes were fallen into 51 clusters, using the 13,821 genes with informative SNPs, which showed expression in 5 DAF endosperm, as the control. The clusters are unevenly distributed on the genome **(Supplemental Figure 3).** Over 34% (86/215) of the rice PEGs were located within the clusters, while less than 16% (27/215) MEGs were distributed in clusters, which suggested that the PEGs tend to be clustered in comparison with the MEGs. Usually, the imprinted genes in the same cluster displayed consistently maternal or paternal expression pattern **(Supplemental Table 4).** This finding implied that there are likely common imprinting regulatory elements to control genes’ imprinting within the same cluster.

### Expression patterns of imprinted genes

We used Genevestigator to study gene expression of the imprinted candidates in rice. The result showed that more than half of the common MEGs of RL-LR and WY-YW are preferentially expressed in the endosperm **(Figure 4A).** Actually, if using the MEGs shared by RL-LR, WY-YW and N9-9N for analysis, the expression preference became more obvious **(Supplemental Figure 4A).** The endosperm preferential expression pattern of MEGs can be observed in *Arabidopsis*, maize, and sorghum (Wolff et al., 2011; Waters et al., 2013; Klosinska et al., 2016; Zhang et al., 2016); therefore, we inferred that MEGs may play an important role in endosperm development. Overall, the PEGs of rice did not display preferential expression in endosperm **(Figure 4A and Supplemental Figure 4B).** Dynamic expression analysis of the imprinted genes during early seed development indicated that many MEGs and some PEGs were not expressed at the first 2–3 days after fertilization **(Figure 4B, Supplemental Figure 4C and D).** Their expression was substantially activated at 3-4 DAF in the endosperm, but usually not in the embryo **(Supplemental Figure 4C).**

**Figure 4.**
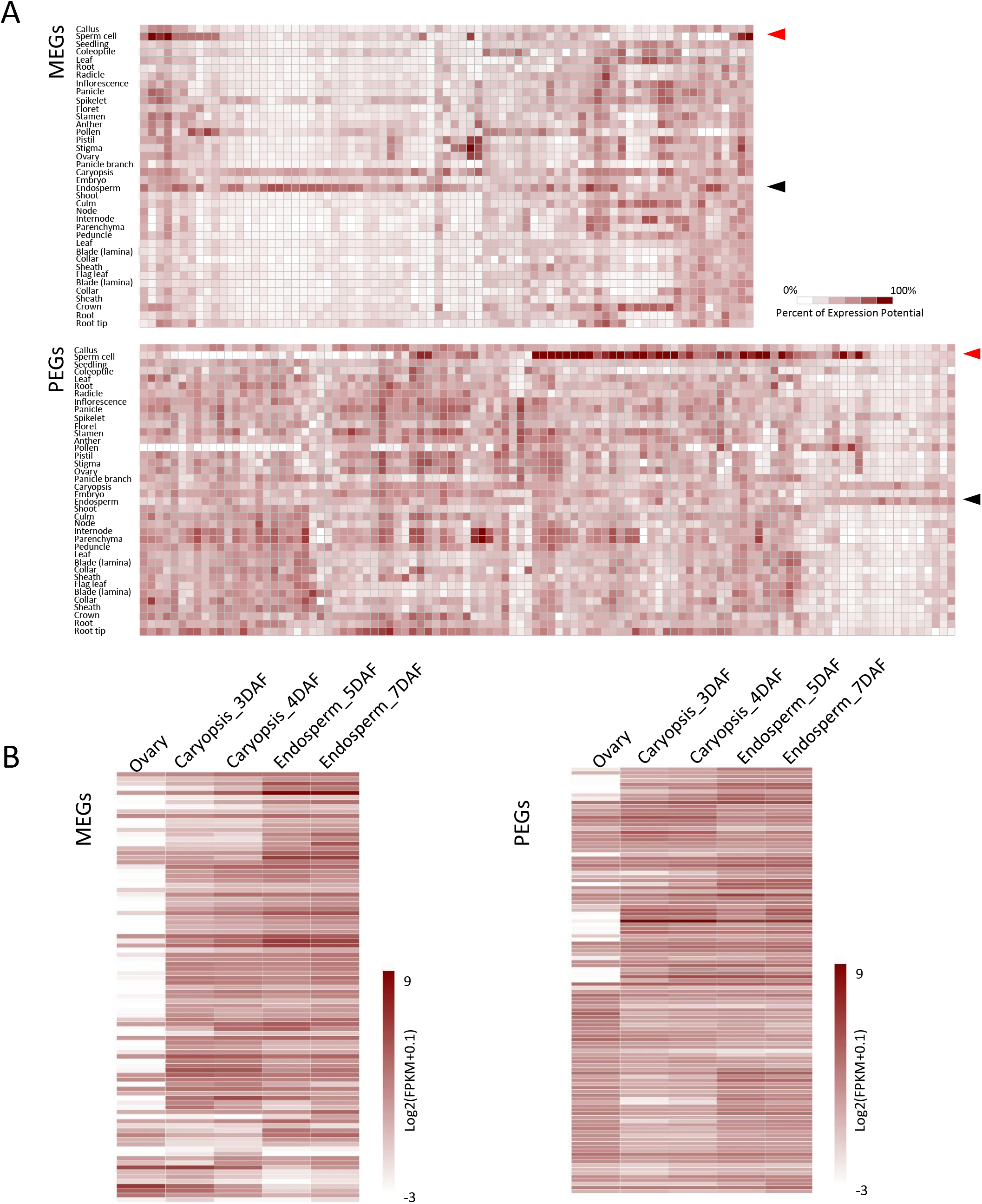
Expression profiles of imprinted genes in rice. **(A)** Expression analysis of the common MEGs (upper panel) and PEGs (lower panel) of RL-LR and WY-YW by Genevestigation. Red and black arrows indicate sperm cell and endosperm, respectively. **(B)** Dynamic expression pattern of the MEGs (left panel) and PEGs (right panel) in developing seed of Nipponbare.

As we learned from *Arabidopsis*, the MEGs are usually activated in the central cell before fertilization (Huh et al., 2008). However, we noted that only approximately half of the MEGs were expressed in the ovary, whereas the rest were predominantly activated in the endosperm after fertilization **(Figure 4A, Supplemental Figure 5A and B).** The finding suggested that some MEGs were likely activated in the central cell, but there are still some other mechanisms to asymmetrically erase epigenetic marks of parental alleles after fertilization; or possibly, erasure of an epigenetic mark in the central cell is not sufficient to trigger gene expression, but requires an activator that exclusively expresses after fertilization. Similarly, approximately half of the PEGs showed the highest expression in sperm cells, but still many did not show expression in sperm cells **(Figure 4A, Supplemental Figure 5A),** which implied that paternal alleles of these PEGs could be specifically activated after fertilization. By surveying publicly available transcriptome data, we found that the expression level of MEGs was generally low in rice sperm cells **(Supplemental Figure 5C).** However, some MEGs were highly expressed in sperm cells **(Supplemental Figure 5A, D and E),** which implied that silencing of the paternal alleles of certain MEGs occurs post fertilization. The findings suggested that regulation of genomic imprinting in rice could be very diverse. Silencing or activation of the paternal or maternal alleles of imprinted genes may occur either pre- or post-fertilization. Moreover, establishing genomic imprinting after fertilization may be popular in rice.

### Epigenetic regulation of gene imprinting in rice

By reanalyzing the publicly available rice DNA methylome data of Nipponbare (Zemach et al., 2010), we found that the DNA methylation profiles of the imprinted genes (RL-LR and YW-WY commons) were quite similar in embryo, root and shoot; however, it was hypomethylated in endosperm **(Supplemental Figure 6).** This mirrored the lower extent of methylation in endosperm on a genome-wide scale (Zemach et al., 2010). In terms of the promoter region, imprinted genes generally exhibited higher CG and CHG methylation levels than their non-imprinted paralogs in the embryo **(Figure 5A and C).** Moreover, MEGs had higher CG and CHG methylation levels than PEGs **(Figure 5A and C).** In the endosperm, CG methylation of the promoter regions of imprinted genes was substantially decreased, while that of the non-imprinted homologous genes was sustained to a resemble extent to the embryo **(Figure 5B).** Notably, MEGs of rice showed higher methylation than that of PEGs in endosperm, which is contrast to the findings in *Arabidopsis* and maize (Pignatta et al., 2014; Zhang et al., 2014). However, this is in agreement with our observation that the rice MEGs showed more enrichment of TEs in the promoter regions in comparison with the PEGs **(Figure 3).** Promoter CHG methylation of the imprinted genes was comparable to their corresponding non-imprinted homologs in the endosperm **(Figure 5D).** The CHH methylation level in the endosperm was very low, and there was no difference between the imprinted genes and their homologs **(Figure 5F).** Distinct to the promoter regions, in the embryo, the gene bodies of PEGs showed higher CG methylation than MEGs **(Figure 5A).** This is consistent with the finding that CG methylation of gene body correlates with transcriptional activity (Zilberman et al., 2007) and the PEGs are overall wildly expressed **(Figure 4A).** However, the higher CG methylation of PEGs in gene bodies was not observable in the endosperm **(Figure 5B).** The results are in agreement with the findings that PEGs are enriched for differential methylated regions (DMRs) in gene bodies, while MEGs tend to show DMRs in promoter regions (Rodrigues et al., 2013). In addition, MEGs showed a higher CHG methylation level of gene body than non-imprinted MEG homologs and PEGs in the embryo **(Figure 5C).** The difference between MEGs and their non-imprinted homologs was not observed in the endosperm, though MEGs were still highly methylated relative to PEGs **(Figure 5D).** Intraspecific variations of methylation can lead to allele-specific imprinting in plants (Pignatta et al., 2014). The methylomic data we used here was collected from Nipponbare (Zemach et al., 2010). To eliminate the genotype impact, we used the highly conserved imprinted genes (22 PEGs and 32 MEGs that shared by all the 4 sets of reciprocals) to investigate the effects of methylation on genomic imprinting. We found that the methylation patterns of the conserved PEGs and MEGs were generally resembled to the common imprinted genes of RL-LR and MY-YM **(Supplemental Figure 7).** Notably, the methylation profiles of the conserved PEGs and their non-imprinted homologs were very alike, either in embryo or endosperm, suggesting that methylation alone likely has quite limited effects on imprinting of conserved PEGs **(Supplemental Figure 8).**

**Figure 5.**
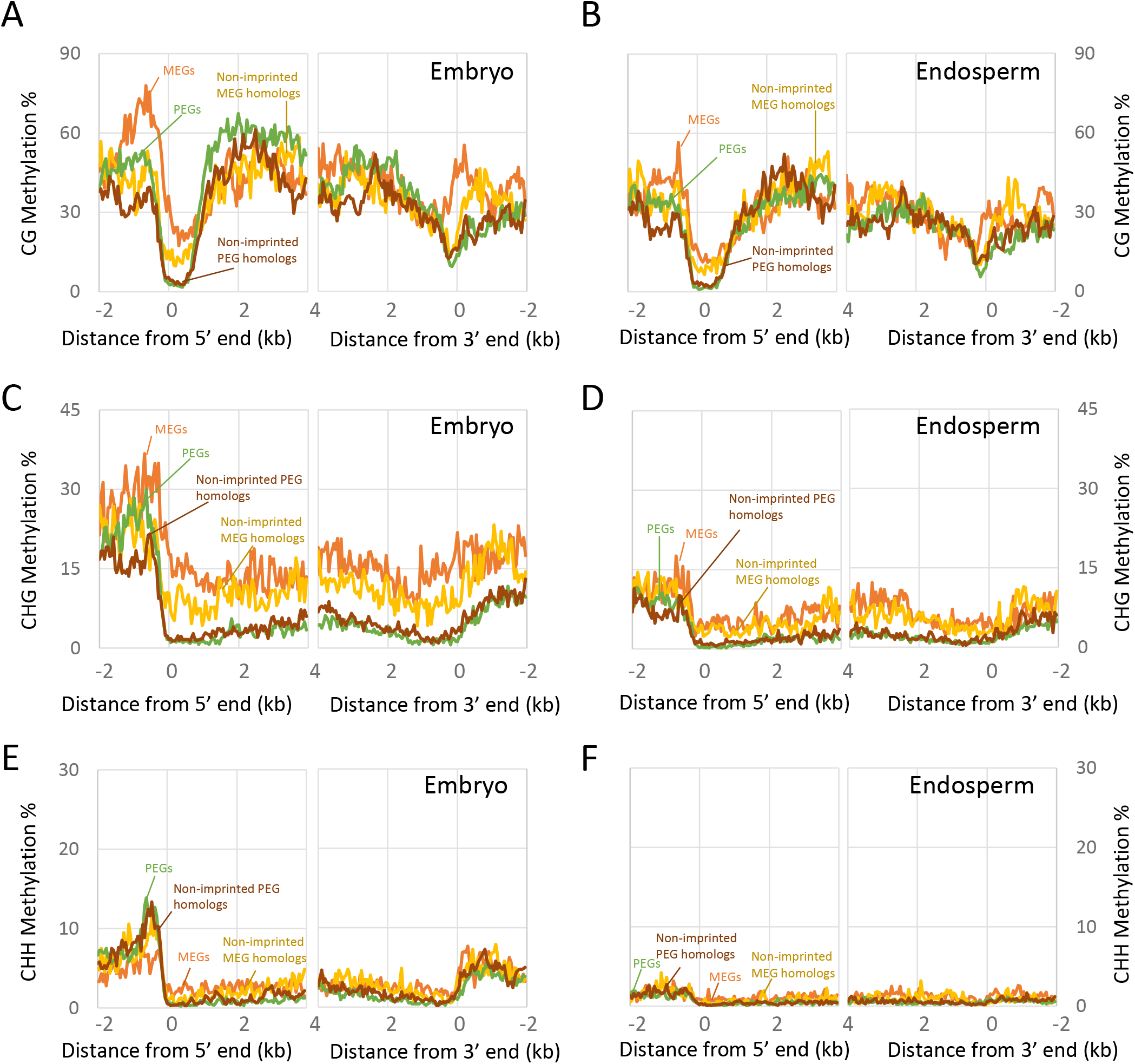
Epigenetic regulation of imprinted genes in rice. The average CG **(A, B)**, CHG **(C, D)** and CHH **(E, F)** methylation profiles of the imprinted genes and their most similar non-imprinted homologs in embryo **(A, C, D)** and endosperm **(B, D, F)**.

Many MEGs do not express in tissues other than the hypomethylated endosperm, while the PEGs are wildly expressed and the imprinting of which shows limited association to DNA methylation. Therefore, we hypothesized that repression of DNA methylation may induce ectopic expression of some MEGs. To test this idea, we applied 5-Aza-2’-Deoxycytidine (AZA), a DNA methylation inhibitor, to the seedlings of Kitaake (*O.sativa* spp. *japonica*) for 10 days. Genome-wide transcriptome analysis showed that the expression of many RL-LR and WY-YW imprinted overlaps was disturbed, and as expected, the MEGs responded to AZA differently from the PEGs **(Supplemental Figure 8A).** The vast majority of PEGs exhibited few expression changes, while approximately 60% of MEGs showed at least 2-fold up-regulation **(Supplemental Figure 8A).** We also noticed that many of the activated MEGs were not expressed in the seedlings under normal conditions **(Supplemental Figure 8B),** implying that decreased methylation did cause ectopic expression of some MEGs. These findings further confirmed that imprinting of the MEGs rely on DNA methylation more than the PEGs.

Generally, the imprinted genes with high expression in seedlings were not sensitive to AZA **(Supplemental Figure 8C).** We assumed that imprinting regulation of these genes may also rely on PRC2-mideated histone modification or some other mechanisms, such as chromatin structure or organization modulating. Malone et al. (2013) profiled H3K27me3 histone modification in young 6-7 DAF rice endosperm. Using this data, we found that approximately 30% (34 out of 115) of the PEGs common in RL-LR and WY-YW coincided with H3K27me3 peaks within the gene body or in the 2-kb flanking regions, whereas only 17% (16 out of 93) of MEGs coincided with H3K27me3 peaks **(Supplemental Table 5).** In agreement with the findings in *Arabidopsis* and maize (Zhang et al., 2014; Moreno-Romero et al., 2016), the results also suggested that the imprinting regulation of many PEGs likely requires both H3K27me3 modification and DNA methylation, whereas that of MEGs relies more on DNA methylation.

### Regulation of genomic imprinting by imprinted small RNAs

Previous studies suggested that small RNAs may involve imprinting regulation (Rodrigues et al., 2013; Pignatta et al., 2014; Yuan et al., 2017). Several dozens of imprinted sRNAs have been found in rice (Rodrigues et al., 2013; Yuan et al., 2017). More than 36% (28/77) of the imprinted sRNA loci are co-localized with the imprinted genes of rice **(Supplemental Table 6).** Interestingly, with no exception, the bias direction of an imprinted 24-nt sRNA loci was opposite to its neighbor imprinted genes **(Supplemental Table 6),** suggesting a maternally expressed 24-nt sRNAs may directly target the paternal allele of its adjacent gene and repress expression, and *vice versa*. To confirm this notion, we investigated the parental expression of all the genes which harbor or adjacent to an imprinted 24-nt sRNA. Forty-two out of the 93 genes have informative SNPs and showed expression in the endosperm. Our results showed that the paternal-sRNA neighbors usually exhibited maternal expression bias (more than 80% transcripts were maternal), while the maternal-sRNA neighbor genes displayed paternal expression bias (less than 50% transcripts were maternal) **(Figure 6A-C).** The results strongly suggested that an imprinted 24-nt sRNAs is able to oppositely target its genic vicinity for imprinting.

**Figure 6.**
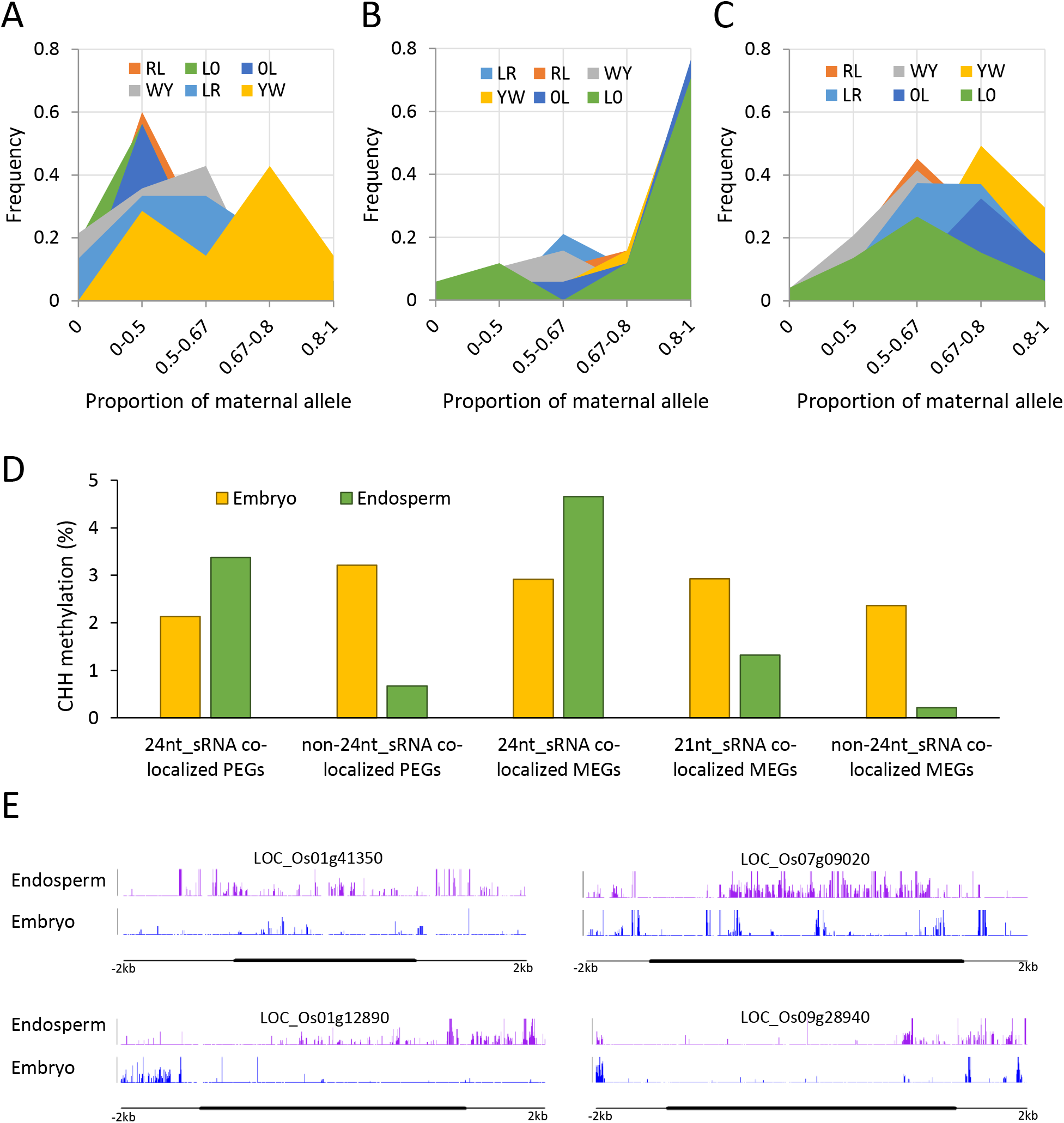
The genic neighbors of imprinted 24-nt sRNAs show opposite parental expression bias and high CHH methylation in endosperm. **(A)** Distribution frequency of the proportion of maternal alleles (FPKM_maternal_/(FPKM_maternal_+FPKM_paternal_)) of the genes that link with a paternally expressed 24-nt sRNA in different crosses. **(B)** Distribution frequency of the proportion of maternal alleles of the genes that link with a paternally expressed 24-nt sRNA in different crosses. **(C)** Distribution frequency of the proportion of maternal alleles of all the testable genes that express in the 5 DAF endosperm and harbor informative SNPs to distinguish the parental expression. **(D)** CHH methylation of the imprinted genes co-localized or not co-localized with 24-nt sRNAs and the imprinted genes co-localized with 21-nt sRNAs in embryo and endosperm. **(E)** Examples of CHH methylation increased imprinted genes that linked with 24-nt sRNAs.

We next compared the CHH methylation level between 24-nt sRNA overlapped and non-overlapped imprinted genes, because 24-nt sRNAs are able to promote CHH methylation through RdDM pathway. The MEGs and PEGs around 24nt-sRNAs both showed high CHH methylation level in the endosperm **(Figure 6D and E),** which is opposite to the observation that the imprinted genes overall showed lower methylation level in the endosperm **(Supplemental Figure 6).** The results indicated that imprinted 24-nt sRNAs possibly target their flanking genes for methylation in an allele specific manner. Notably, distinct from the 24-nt sRNAs, all the 5 imprinted 21-nt sRNAs showed the same maternal bias as the MEGs they overlapped **(Supplemental Table 6 and** Yuan et al., 2017), which likely formed imprinted clusters with their vicinities. Moreover, the CHH methylation pattern of the 21nt-sRNAs associated imprinted genes was resemble to the control **(Figure 6D).**

### Domestication shows limited influence on genomic imprinting

The imprinting conservation between cultivated rice and wild rice (*Oryza rufipogon*) remains unknown. We made interspecific crosses between cultivated rice and Dongxiang wild rice (*O. rufipogon*, designated as “D” hereafter). By an RT-PCR sequencing approach, thirteen out of the 14 confirmed imprinted genes of cultivated rice **(Supplemental Figure 1)** also displayed the parent-of-origin expression pattern **(Figure 7),** which suggested that the regulation of genomic imprinting is likely very conserved between *O. sativa* and *O. rufipogon*, and domestication likely imposed limited effects on the evolution of genomic imprinting in rice.

**Figure 7.**
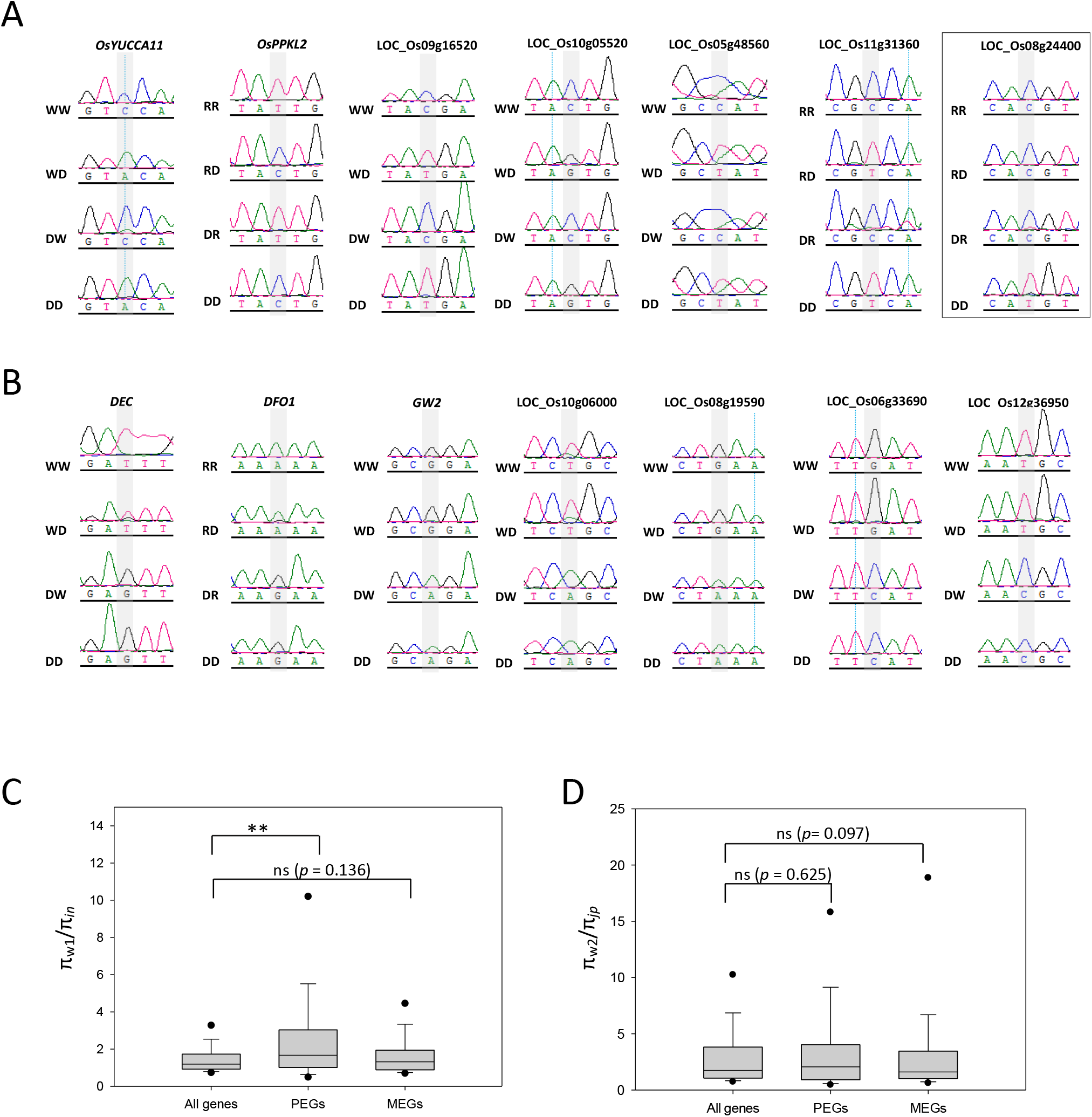
Detection of gene imprinting status in cultivated rice/wild rice reciprocal crosses. **(A, B)** Imprinting status of the PEGs **(A)** and MEGs **(B)** identified from RL-LR and WY-YW in cultivar (*O. sativa*) x wild rice (*O. rufipogon*) reciprocal crosses. Polymorphic sites are shaded in gray. W, R and D indicate Wufeng, Rongfeng and Dongxiang wild rice, respectively. The gene that does not display a parent-of-origin expression pattern is boxed. **(C, D)** Difference in nucleotide diversity (π) between Wl-type wild rice and indica **(C)** and between W2-type wild rice and japonica **(D)**. Mann–Whitney rank sum test is used for the statistical analysis. ** indicates *p*<0.01. All genes encoded by the rice genome are used as the control.

We further investigated the demographic history of imprinted loci. Recent evidence suggests that W1 type wild rice is phylogenetically closer to *indica*, whereas the W2 type is closer to *japonica* rice (Huang et al., 2012; Xu et al., 2012). Here we used π_w1_/π_*in*_ and π_w2_/π_*ja*_ to indicate the difference in nucleotide diversity between wild rice and *indica* or *japonica*. In *indica*, the π_w1_/π_*in*_ of PEGs was significantly higher than that of the whole genome (*p* < 0.001, **Figure 7C).** The reduced nucleotide diversity of PEGs in *indica* indicates that possibly the genes underwent selection during domestication. However, for *japonica*, the imprinted genes showed no evidence of being selected **(Figure 7D).** We did not find the π_w1_/π_*in*_ or π_w2_/π_*ja*_ of MEGs was significantly different from that of *indica* and *japonica* genome **(Figure 10C and D).** Taken together, we inferred that most of the imprinted genes were not extensively selected during domestication.

**Genomic imprinting is conserved at some loci in plants**

We searched the imprinted genes identified from other plant species (Gehring et al., 2011; Schoft et al., 2011; Waters et al., 2011; Wolff et al., 2011; Zhang et al., 2011; Waters et al., 2013; Pignatta et al., 2014; Zhang et al., 2014) against the rice genome to obtain their rice orthologs. If just considering the top hits, we found 7, 53 and 32 imprinted genes of *Arabidopsis*, maize and sorghum, respectively, sharing imprinted orthologs in rice **(Figure 8A, Supplemental Table 7),** which indicated that about 13% (78/588) imprinted genes in rice may also be imprinted in other plant species. When using the top 3 hits, the number increased to 24% (143/589) **(Supplemental Table 7).** The result confirmed relatively low conservation of genomic imprinting in plants (Waters et al., 2013; Hatorangan et al., 2016). However, we found some conserved imprinted loci among plant species **(Supplemental Table 7).** To test the conservation, we constructed reciprocal crosses of barley, using the varieties Morex and Bowman as the parents. First, we searched the most similar barley orthologs of some conserved imprinted genes **(Supplemental Table 7),** using the rice protein sequences as the queries. By searching the variation information deposited in the Gramene database (ensembl.gramene.org/Hordeum_vulgare/lnfo/lndex), we designed primers to amplify cDNA fragments with potentially informative SNPs. Among the 21 genes we tested, 10 lacked informative SNPs or expression in the endosperm **(Figure 8B).** Of the remaining 11 genes, two were identified as MEGs and 6 as PEGs **(Figure 8C).** In addition, one showed odd monoallelic expression in a parent-of-origin-independent manner, and 2 were biallelic **(Figure 8C).** In total, approximately 73% (8 out of 11) of the determinable genes showed parent-of-origin expression patterns, which confirmed the imprinting conservation at some loci in plants, at least in monocots.

**Figure 8.**
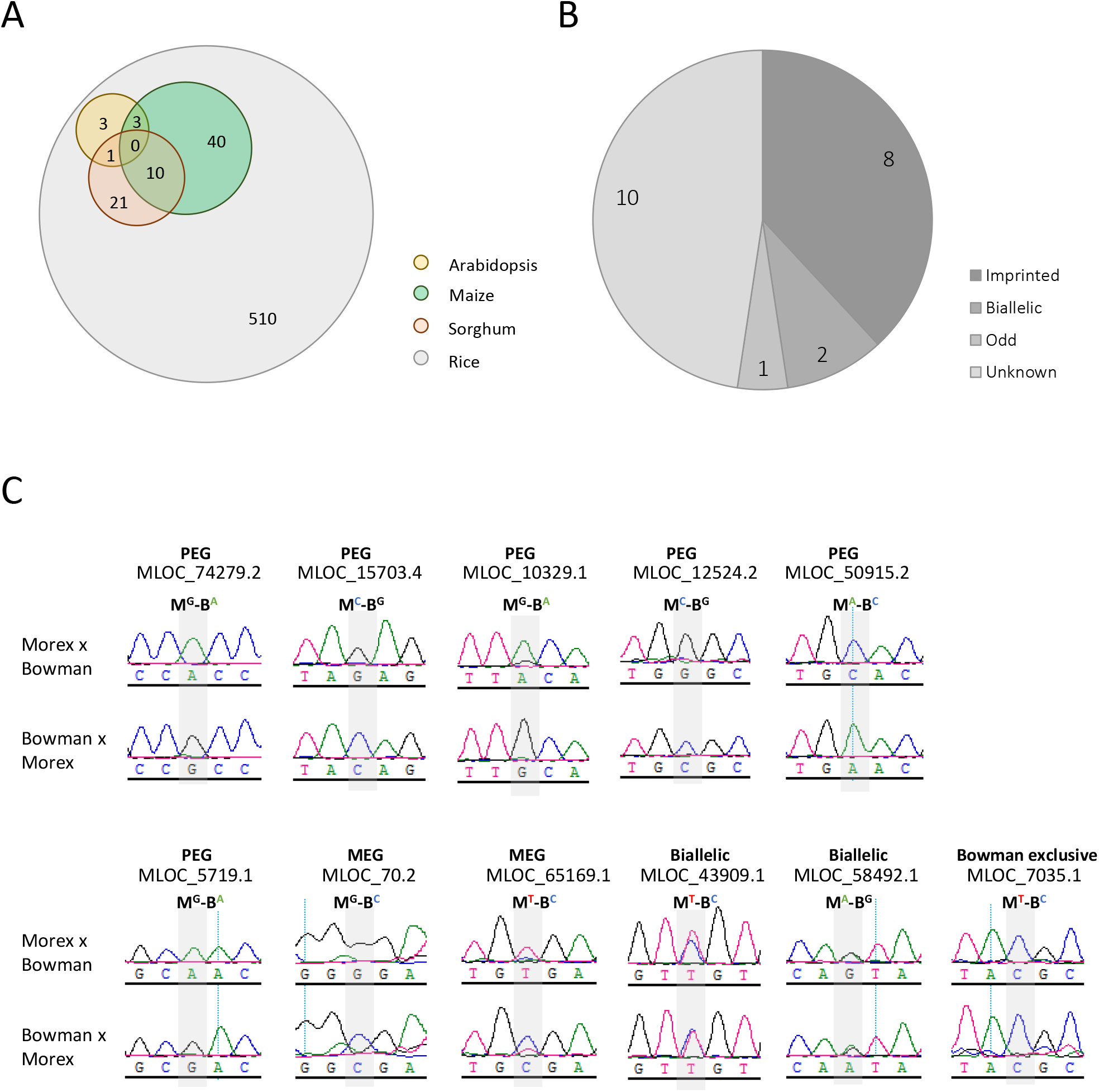
Some loci exhibited conserved imprinting in plants. **(A)** Veen diagram analysis of the imprinting conservation in plants. Only a few of the most similar rice homologs (top 1 hits) of the imprinted genes that found in maize, sorghum and *Arabidopsis* show imprinting in rice. **(B)** Many of the conserved imprinting loci are imprinted in barley. Conserved imprinted genes here refer to the ones identified from at least two species of rice, maize, sorghum and *Arabidopsis*. The top 3 blast hits (e-value < 10e-10) are used to evaluate the imprinting conservation. **(C)** Validation of the imprinting status of 11 conserved imprinted loci in barley. M and B indicate the varieties Morex and Bowman, respectively, and the superscripts indicate the polymorphic nucleotides of each allele. Polymorphic sites are highlighted in gray.

## Discussion

By generating reciprocal crosses of *indica* and *japonica* rice varieties, here we identified 208 strong imprinted genes with 5-fold higher than the 2:1 ratio that shared by the RL-LR and WY-YW reciprocal-cross sets **(Supplemental Table 1)**. Thirteen out of the 14 imprinted candidates genes identified from cultivated rice also showed parent-of-origin expression in the *O. sativa* and *O. rufipogon* interspecific reciprocal crosses **(Figure 7A and B),** indicating that the regulation of genomic imprinting of the two rice species are likely very conserved, and wild rice possibly has similar genomic imprinting as the cultivars. The process of rice domestication likely imposed limited force on the evolution of imprinting, though PEGs were tended to be selected during *indica* rice domestication **(Figure 7C).** Klosinska et al.(2016) found that the parentally biased expression pattern is conserved between *Arabidopsis lyrata* and *A.thaliana*. These findings suggested that there is evolutionary conservation in imprinting between species with a close phylogenetic relationship. In the present study, we comprehensively compared the rice imprinted genes to those identified from other plant species. We found that about 13-24% of the rice imprinted genes had imprinted homologs at least in two species **(Figure 8A and Supplemental Table 7)**. This is in agreement with previously estimation that about 20% imprinting is conserved between species (Waters et al., 2013; Hatorangan et al., 2016). However, we here found some loci are tend to be imprinted in different plant species **(Figure 8).** Auxin is important for triggering central cell division and for maintaining endosperm development (Bernardi et al., 2012; Figueiredo et al., 2015). *OsYUCCA11* of rice, *ZmYUCCA1* of maize and *AtYUCCA10* of *Arabidopsis* are orthologs, and both of which were identified as PEGs **(Supplemental Table 7).** *Tryptophan aminotransferase related 1 (TAR1*), another auxin biosynthetic gene, was identified as a PEG in different species **(Supplemental Table 7).** *OsTAR1* and *OsYUCCA11* contribute to auxin accumulation during rice grain development (Abu-Zaitoon et al., 2012). Likewise, *ZmYUCCA1* is important for endosperm development of maize (Bernardi et al., 2012). These findings indicate that genomic imprinting may have evolutionary significance and directly function in auxin-mediated seed development. Xin et al. (2013) also found that many auxin-related genes showed expression bias in maize. In addition, we noted that several imprinted genes involving epigenetic regulation, such as chromatin-remodeling factors, histone H3K9 methyltransferase genes, *Variant In Methylation* gene, were identified as PEGs in different plant species **(Supplemental Table 7).** For example, *Arabidopsis SUVH7* encodes a SET domain protein and acts as a histone methyltransferase for epigenetic control of gene expression, which is essential to establish postzygotic hybridization barriers (Wolff et al., 2015). Imprinting is conserved in its orthologs of rice and maize, which led us to assume that these genes may contribute to reproductive isolation in both monocots and dicots. We randomly chose some imprinted genes that at least identified from two species, to detect the imprinting status of their most similar homologs in barley. More than 70% of the genes we tested showed parent-of-origin expression pattern **(Figure 8B),** which convinced us that there is imprinting conservation at some loci in plants, at least in monocots.

The studies of *Brassicaceae* species revealed that many imprinted genes were often physically located in the vicinity of TEs (Wolff et al., 2011; Pignatta et al., 2014; Hatorangan et al., 2016). TEs were found to be activated in central cells as well as in the vegetative cells of pollen (Slotkin et al., 2009; Schoft et al., 2011; Calarco et al., 2012; Ibarra et al., 2012). Genomic imprinting was believed to associate with the dynamic regulation of TEs in the reproductive cells and endosperm (Gehring et al., 2009). Asymmetrical activation of TEs in the central and sperm cell leads to monoallelic expression after fertilization. However, our study suggested that the TEs are not enriched around the imprinted genes of rice **(Figure 3).** Interestingly, rice PEGs showed less association to the TEs than the MEGs **(Figure 3),** which is opposite to the findings in *Arabidopsis* (Pignatta et al., 2014). Unlike the gene silencing observed in *Arabidopsis*, many TEs are not silenced in the sperm cells of rice (Anderson et al., 2013). We assumed that the regulation of imprinting in rice may be different.

Many MEGs are activated before fertilization in *Arabidopsis*. For example, the PRC2 components *MEA* and *FIS2* are highly expressed in central cells and gradually repressed after fertilization(Luo et al., 2000; Baroux et al., 2006). Hypomethylated environment of the central cell (Gehring et al., 2006; Jullien et al., 2006; Park et al., 2016) promotes maternal allele expression in the endosperm, whereas the paternal allele remains hypermethylated and silenced in the sperm cell and endosperm. Per this model, MEG expression persists immediately after fertilization. However, in contrast, we found many MEGs of rice to be specifically activated at 3–4 DAF in the endosperm **(Figure 4B, Supplemental Figure 4C and D).** As an example, PRC2 component gene *OsFIE1* is an exclusively endosperm-expressed gene (Zhang et al., 2012a). *OsFIE1* was not detectable before fertilization until 3 DAF **(Supplemental Figure 4D)**. Though we were short of evidence to show that *OsFIE1* was absent in central cells, the findings indicated that imprinting regulation of *OsFIE1* is distinct from that of *MEA* and *FIS2* in *Arabidopsis*. Extensive activation of imprinted genes after fertilization was also observed in maize (Xin et al., 2013). *Arabidopsis* MEG *AtFH5* was specifically activated in the endosperm as well (Gerald et al., 2009). These findings suggested that for some MEGs, differential imprinting between two parental alleles is likely expressed in endosperm but not in germ cells. Xing et al. (2015) found a dramatic decrease in global DNA methylation in the endosperm of rice from 2–3 DAF. The methylation level of endosperm is even lower than that of the central cell (Park et al., 2016). Imprinting of these genes could be caused by asymmetric DNA demethylation in the endosperm. We also noticed that several MEGs were highly expressed in sperm cells **(Supplemental Figure 5),** but their paternal alleles were silenced after fertilization, which indicates that plants can selectively apply imprinting markers to the paternal genome to repress gene expression after fertilization. However, the underlying mechanism still awaits elucidation. Imprinting regulation of PEGs is poorly understood so far. We found that >50% of PEGs were highly expressed in sperm cells **(Figure 4A).** Anderson et al.(Anderson et al., 2013) showed low expression of *MET1* and high expression of *ROS1a*, a DNA glycosylase reversing DNA methylation, in rice sperm cells. The results suggested that possibly, similar to what happens in central cells, the hypomethylated environment in the sperm cell facilitates paternal monoallelic expression of PEGs in the endosperm. The RdDM pathway is considered to regulate paternal genomic imprinting in *Arabidopsis* (Vu et al., 2013). However, by surveying the transcriptome, Anderson et al. (2013) suggested that evidence of RNAi activity in sperm cells of rice is lacking. Nonetheless, our result indicated that the 24-nt small RNAs may be involved in the regulation of genomic imprinting in rice **(Figure 6).** The imprinted sRNAs’ genic vicinities were generally showed opposite parental expression pattern to the imprinted sRNAs **(Figure 6).**

Several imprinted genes we found in rice have been functionally well studied. Among them, *GW2* and *OsPPKL2* are characterized as two quantitative trait loci determining seed size in rice, and they also function in grain filling and starch accumulation (Song et al., 2007; Zhang et al., 2012b). *OsCBL2* is likely involved in the development of aleurone layer (Hwang et al., 2005). Notably, *Os8N3*, which shows the ability of copper reallocation, was initially identified as a disease-resistance gene, then found to be essential for pollen and seed development (Yang et al., 2006; Yuan et al., 2010). The *CAS* gene triggers expression of sugar partitioning and metabolic genes during pollen and seed development (Zhang et al., 2010). These genes are likely involved in the reallocation of resources and nutrients from maternal tissues to offspring, which provide new support for the kinship theory and suggest that at least some imprinted genes are indispensable for proper seed development. Interestingly, activation of many imprinted genes in the endosperm at 3–4 DAF **(Figure 4B and Supplemental Figure 4)** coincides with the cellularization event in rice (Ishikawa et al., 2011; Chen et al., 2016). Genetic evidence showed that *OsMADS87*, a transiently activated MEG, can promote endosperm proliferation and repress its cellularization (Ishikawa et al., 2011; Chen et al., 2016). In contrast, the MEG *OsFIE1* promotes cellularization (Folsom et al., 2014). Collectively, we believe that many imprinted genes in rice may have biological significance for rice seed development (Yuan et al., 2017).

## Methods

### Plant materials and growth conditions

Wufeng-A and -B, Rongfeng-A and -B, Liuqianxin-A and -B, Yu6-A and -B, and wild rice were grown in Nanchang, Jiangxi Province, China. Wild rice is an often-outcrossing plant; therefore, to avoid variation within the genome, Dongxiang wild rice was self-fertilized by bagging for six generations and then used for crossing. The RL-LR and WY-YW reciprocal crosses were performed in the summer season of 2014. Plants were grown in a paddy field under regular conditions. The ovary donors were emasculated before flowering and moved into a growth chamber on a 14h/10h light-dark cycle at 30°C. At 5 DAF, the endosperm was carefully collected with a pipette to avoid maternal tissue contamination per the method used by Luo et al. (2011). Two (WY-YW) or three (LR-RL) replicates were set for mining of imprinted genes. Each replicate was pooled with 20–30 endosperm produced by the hybrids. *Japonica* varieties Nipponbare and Kitaake were grown in a growth chamber (14h/10h light-dark cycle, 30°C) and labeled when flowering. The caryopsis at 0 to 3 DAF or endosperm at 5 to 7 DAF were collected for RNA isolation. Barley varieties Morex and Bowman were planted in the experimental plot with regular nutrient and water management in Chengdu, Sichuan Province, China. The reciprocal crosses were performed in the field. The ovary donors were emasculated before flowering. At 10 DAF, the endosperm of the hybrid seeds was collected with care.

For the 5-aza-2’-deoxycytidine treatment assay, Kitaake seeds were sterilized and grown on 1/2 MS medium with 70 mg.L^−1^ AZA for 10 days in a growth chamber at a constant temperature of 28°C with a 14h/10h light-dark cycle. The controls were grown under the same conditions, except the medium was AZA free. Seedlings were collected for RNA-seq. Three biological replicates were set, each one comprised approximately 10 individuals.

### DNA and RNA isolation and Real-Time PCR assay

DNA was extracted from 100 mg leaves of Liuqianxin-A, Yu6-A, Rongfeng-B, and Wufeng-B plants using the EasyPure Plant Genomic DNA Kit (TRANS) following the manufacturer’s suggested procedures. RNA was extracted from endosperm or seedlings using a Plant RNA Kit (OMEGA) and treated with RNA-free DNase set (OMEGA) to discard DNA contamination per the manufacturer’s protocol.

Complementary DNA was synthesized using the PrimeScript RT Master Mix (TAKARA). 2 μL of the 5x diluted complementary DNA was used for qPCR in a 20-μL reaction using AceQ^®^ qPCR SYBR^®^ Green Master Mix (Vazyme). The reactions were performed on the CFX Connect^™^ Real-Time System (BioRad). The proteasome gene (LOC_Os03g63430) was used as the endogenous control. For each sample, we performed at least three independent biological replicates with three technical replicates. The Relative expression level was calculated by the ΔΔCt approach. All the primers used in the present study can be found in **Supplemental Table 8**.

### DNA re-sequencing, RNA-seq, and imprinted gene identification

Shredded DNA was used for library preparation following the protocols of NEBNext Ultra DNA Library Prep Kit for Illumina. The libraries were qualified by Qubit^®^ 2.0 Fluorometer and Agilent Bioanalyzer 2100 (Agilent Technologies) and then sequenced on the Illumina HiSeq platform. Purified RNA was inspected the integrity by Agilent Bioanalyzer 2100 (Agilent Technologies). Qualified total RNA was used for RNA-seq library preparation with the TruSeq RNA Kit v2 (Illumina). Libraries were sequenced on the Illumina HiSeq platform.

Raw reads were processed by the package of Trimmomatic v0.32 to remove the adaptors and discard low quality reads. The filtered reads then were aligned to the rice reference genome (MSU v7.0, rice.plantbiology.msu.edu) using bowtie2. Two mismatches were allowed for alignment. Reads that mapped to multiple sites were discarded. GATK was used to process aligned reads for SNPs calling. The obtained SNPs were then filtered with genome coverage, and the ones with depths <11 were removed. RNA alignment was carried out by bowtie2. Then, samtllos (version 0.1.16) was used to process the aligned reads and call SNPs with default parameters. RNA SNPs were aligned to DNA SNPs with in-house Perl script of which depths <11 were removed. Reads that could align to multiple loci were discarded. Imprinted genes were identified by calculating the possibility of (maternal reads):(paternal reads) at each SNP site deviating from the expected 2:1 ratio using *χ*^2^ tests.

### Validation of imprinted genes

Primers were designed to amplify 300–500-bp gene fragments harboring informative SNPs from the cDNA of reciprocal hybrids and of the parents. The PCR products were purified for Sanger sequencing or cloned into pEAZY-Blunt Zero (TRANS) for the single-colony sequencing assay; at least 12 colonies were sequenced. For the validation of *GW2* and *OsPPKL2* by the pyro-sequencing approach, genes were amplified with 2x EPIK Amplification Mix (Bioline) using cDNA as the template. Fifteen to 20-μL PCR products were sequenced on PyroMark Q96 (Qiagen) platform. The primers used for the assays are listed in **Supplemental Table 8.**

### Epigenetic analysis of imprinted genes

Rice bisulfite-sequencing data of root, shoot, embryo, and endosperm was obtained from the NCBI GEO database (Accession: GSE22591). The reads were aligned to the rice reference genome using Bismark Extractor software, allowing two mismatches per read. Reads that mapped to multiple positions were discarded. Then, the average methylation level was calculated at each cytosine in the context of CG, CHG, and CHH of the PEGs, MEGs, MEG homologs, and PEG homologs, and visualized in each 50 bp bin.

Data on H3K27me3 in rice endosperm was obtained from the NCBI GEO database (Accession: GSE27048). Reads were aligned to the rice genome using Bowtie2. Two mismatches per read were allowed, and reads that mapped to more than one site were removed. The program MACS was used to process aligned reads and call peaks as described in Malone et al. (2011). Peaks with fold changes >5 in comparison with the mock and *q* value < 0.05 were kept for further analysis.

### Characterization of imprinted genes

The information on imprinted small RNAs in rice endosperm was obtained from previous studies (Rodrigues et al., 2013; Yuan et al., 2017). Gene expression data was obtained from Genevestigator, RiceXPro (http://ricexpro.dna.affrc.go.jp) and NCBI Gene Expression Omnibus (GEO, Accession: GSE50777 and GSE40674). The expression profiles of the imprinted genes in different tissues were generated by Genevestigator, using all the expression dataset of wild-type rice that collected by Genevestigator. We used the transcriptome data of early development seeds of rice that deposited in RiceXPro (http://ricexpro.dna.affrc.go.jp) to analysis the expression pattern of the shared imprinted genes among RL-LR, WY-YW and N9-9N. Venn diagram analysis was performed on the web-based tool VENNY 2.1 (http://bioinfogp.cnb.csic.es/tools/venny/index.html). Gene Ontology analysis was performed by the online tool of Singular Enrichment Analysis (SEA) provided by AgriGO (http://bioinfo.cau.edu.cn/agriGO/). Gene clustering analysis was conducted by the software REEF (Coppe et al., 2006) with window size 100 kb, step size 50 kb and default statistical settings. The genes expressed in the endosperm at 5 DAF which showed polymorphisms between the parents of RL-LR and WY-YW were used as the reference for the GO analysis and the clustering analysis. For TEs annotation, we downloaded all of the transposable element families in rice (http://genome.arizona.edu/cgi-bin/rite/index.cgi) as the source for RepeatModeler modeling, RepeatModeler will identify repeat element boundaries and family relationships from those transposable element sequences. Then, we combine the database with the Dfam database as the combined database, and use RepeatMasker to screen DNA sequences for transposable elements in Nipponbase.

### Differential expression analysis

The raw reads were filtered by Seqtk before mapping to genome using Tophat (version: 2.0.9). The fragments of genes were counted using HTSeq followed by TMM (trimmed mean of M values) normalization. Significant differential expressed genes (DEGs) were identified as those with a False Discovery Rate (FDR) value above the threshold (Q< 0.05) and fold-change >2 using edgeR software.

### Evolutionary analysis of imprinted genes

We used the published resequencing data of 50 *japonica* and 25 *O. rufipogon* (Xu et al., 2012) to calculate the polymorphism across all W1 type wild rice and *indica*, and W2 type wild rice and *japonica* to test selection of the imprinted genes in rice. The sequence diversity ratio in each 5 kb region with 2.5 kb sliding were calculated using License (version 3.0). Mann–Whitney rank sum test was used to determine the statistical significance by SigmaPlot 12.5.

## Author contributions

C.C., Z.E. and Q.Q.L designed the research. C.C. wrote the manuscript. T.L., S.Z., Z.L., Z.S. and S.L. performed the research. X.Z, R.C., J.H., Y.S. and L.W. analyzed the data. All authors commented on the manuscript and contributed to the writing.

## Acknowledgments

This research was supported by grants from National Key Research and Development Program of China (2016YFD0100902 to C.C.), Natural Science Foundation of Jiangsu Province (BK20150446 to C.C), National Natural Science Foundation of China (31571623 to C.C and 31671964 to R.C.) and the Priority Academic Development of Jiangsu Higher Education Institutions.

No conflict of interest declared.

## Supplemental Data

**Supplemental Figure** 1. Validation of the imprinting status of selected candidates.

**Supplemental Figure 2.**. Heat map of the proportion of maternal alleles of all the imprinted genes in different cross combinations.

**Supplemental Figure 3.** Distribution of imprinted clusters in the rice genome.

**Supplemental Figure 4.** Expression profiles of the high-convincing imprinted candidates (RL-LR, WY-YW and N9-9N commons) and their most similar homologs in rice.

**Supplemental Figure 5.** Gene expression of the imprinted candidates in reproductive cells or tissues.

**Supplemental Figure 6.** DNA Methylation profiles of the imprinted genes (RL-LR and WY-YW commons) in root, shoot, embryo and endosperm in the context of CG, CHG and CHH in rice.

**Supplemental Figure 7.** DNA Methylation profiles of the imprinted genes (RL-LR, WY-YW, N9-9N and L0-0L commons) and their most similar homologs in embryo and endosperm in context of CG, CHG and CHH of rice.

**Supplemental Figure 8.** Differential responses of PEGs and MEGs to the methylation inhibitor 5-Aza-2’-Deoxycytidine (AZA) in rice seedlings.

**Supplemental Table 1.** Imprinted genes identified in Rice.

**Supplemental Table 2.** Epigenetic and auxin related imprinted genes of rice. **Supplemental Table 3.** Allele-specific imprinting of rice.

**Supplemental Table 4.** Imprinted clusters of rice.

**Supplemental Table 5.** Enrichment of H3K27me3 in the 2-kb flanking regions of imprinted genes in rice endosperm

**Supplemental Table 6.** Imprinted genes adjacent to imprinted small RNAs in rice.

**Supplemental Table 7.** Conservation of some imprinted loci in different plant species.

**Supplemental Table 8.** Primers used in the present study.

